# Real-time monitoring of cell surface protein arrival with split luciferases

**DOI:** 10.1101/2023.02.22.529559

**Authors:** Alexandra A.M. Fischer, Julia Baaske, Winfried Römer, Wilfried Weber, Roland Thuenauer

**Affiliations:** Signaling Research Centres BIOSS and CIBSS and Faculty of Biology, University of Freiburg, Freiburg, Germany; Faculty of Biology, University of Freiburg, 79104 Freiburg, Germany; Spemann Graduate School of Biology and Medicine (SGBM), University of Freiburg; Center for Structural Systems Biology (CSSB), Hamburg, Germany; Technology Platform Light Microscopy, University of Hamburg, Hamburg, Germany; Technology Platform Microscopy and Image Analysis (TP MIA), Leibniz Institute of Virology (LIV), Hamburg, Germany

**Keywords:** Split luciferases, epithelial cells, cell polarization, protein sorting, apical membrane traffic, basolateral membrane traffic

## Abstract

Each cell in a multicellular organism permanently adjusts the concentration of its cell surface proteins. In particular, epithelial cells tightly control the number of carriers, transporters and cell adhesion proteins at their plasma membrane. However, the sensitively measuring the cell surface concentration of a particular protein of interest in live cells and in real time represents a considerable challenge.

Here, we introduce a novel approach based on split luciferases, which uses one luciferase fragment as tag on the protein of interest and the second fragment as supplement to the extracellular medium. Once the protein of interest arrives at the cell surface, the luciferase fragments complement and generate luminescence. We compared the performance of split Gaussia luciferase and split Nanoluciferase by using a system to synchronize biosynthetic trafficking with conditional aggregation domains. The best results were achieved with split Nanoluciferase, for which luminescence increased more than 6000-fold upon recombination. Furthermore, we showed that our approach can separately detect and quantify the arrival of membrane proteins at the apical and basolateral plasma membrane in single polarized epithelial cells by detecting the luminescence signals with a microscope, thus opening novel avenues for characterizing the variations in trafficking in individual epithelial cells.

**Synopsis:** We present here a novel method to determine the arrival of a membrane protein of interest at the cell surface in real time and with sufficient sensitivity to achieve single-cell resolution. This allows measuring detailed kinetics of plasma membrane trafficking and thus to detect also minor changes in protein sorting and intracellular trafficking. Furthermore, the single-cell sensitivity of the method will enable to systematically characterize variations in trafficking between neighboring cells within a multicellular organism.

## Introduction

The cell surface concentration of membrane proteins, such as receptors or cell adhesion proteins, determines how a cell communicates and interacts with its environment and thus ensures the proper function of a cell within a multicellular organism. The localization of membrane proteins at the plasma membrane is regulated by biosynthetic, endocytic, recycling, and degradative trafficking pathways, which enable the cell to adjust the cell surface concentration of specific proteins on a timescale of seconds to minutes [1–3].

Exact monitoring of changes in the cell surface concentration of a membrane protein of interest (POI) in live cells in real time represents a considerable challenge. Direct quantification with fluorescence microscopy is currently not achievable, because whole live cells would have to be repeatedly imaged with a spatial resolution of <5 nm to reliably discern proteins in the plasma membrane from proteins that are still at the cytoplasm, e.g. in endosomes adjacent to the plasma membrane. An approach to resolve this ‘resolution problem’ are membrane impermeant labels for the POI, which specifically bind and therefore label proteins at the cell surface when applied to the extracellular medium. For example, addition of fluorescently labeled antibodies, which bind to the extracellular domain of the POI, represents a commonly used strategy to measure the plasma membrane concentration of proteins in flow cytometry [4] and can also be adapted for adherent cells [5]. Nevertheless, when applying fluorescently labeled antibodies to the extracellular medium, considerable background signal from unbound antibodies prevents a precise and sensitive measurement of the signal from bound antibodies. To avoid this, unbound antibodies can be removed by additional washing steps, which, however, precludes real-time monitoring.

To avoid washing steps, a probe is required that only produces a signal when bound to the POI. An ideal probe for measuring rapid changes in cell surface concentrations of proteins needs to fulfil certain conditions: (1) Signal levels should increase significantly upon binding, ideally by several orders of magnitude, to enable sensitive detection and limiting the required probe concentration. (2) Signals should be generated rapidly after binding, ideally within <1 min, to follow rapid changes in cell surface protein concentration. It is worth noting that rapid and stable signal generation upon binding benefits from high on-rates (k_on_) of the probe and low dissociation constants (k_d_). Two different types of approaches for generating signals only upon cell surface arrival of proteins have been developed. The first type is based on fluorogenic probes, which increase their fluorescence when bound, and the second type is based on split reporters, which only generate signals upon recombination of the split reporter fragments. Nevertheless, all approaches reported so far have limitations, and none simultaneously meets condition (1) and (2) as defined before. This allowed to utilize them only for specific applications: For example, membrane-impermeable fluorogenic probes that can be specifically attached to proteins of interest via SNAP and HALO tags have been recently developed [6]. However, SNAP-based probes have a rather low k_on_ in the range of 10^3^ M^-1^s^-1^ [7], which means that very high concentrations of the probe have to be used and fast kinetic changes cannot be properly monitored. HALO-based probes do have 1000 - 10000 times higher k_on_ values, but available probes do show only a six-fold increase of fluorescence signal levels upon binding [6], which results in comparably high background signals from unbound probes, thus limiting the sensitivity of the assay. Split GFP variants represent the best studied split reporters that have been utilized for measuring the cell surface concentration of proteins [8]. However, recombined split GFPs need typically >15 min to become fully fluorescent [9,10]. This slow kinetics makes it impossible to use split GFPs for measuring changes in cell surface protein concentrations in real time.

Here, we adapt split luciferases for measuring the cell surface concentration of membrane proteins in real time. We tested split Gaussia luciferase (GLuc) [11] and split Nanoluciferase [12] for real-time monitoring of the cell surface arrival of membrane proteins that were synchronously released from the ER using a conditional aggregation domain (CAD)-based system [13]. Recombined split Nanoluciferase yielded much higher signals and was therefore characterized further. Luminescence was detectable within <1 min after recombination of split Nanoluciferase and luminescence levels increased more than 6000-fold upon recombination. The split Nanoluciferase has superior kinetics and signal-increase properties. Furthermore, a small tag of only 11 amino acids needs to be added to the POI, which is available as variants with k_d_ values ranging from 10^−5^ M to 10^−10^ M [12]. All that makes this system a powerful and versatile tool for monitoring cell surface concentrations of proteins in real time. To demonstrate the capabilities of the system we show real-time measurements of cell surface arrival in single cells via microscopy and specific measurements of apical and basolateral arrival in polarized epithelial cells.

## Results

### Functional principle of split luficerase-mediated detection of cell surface arrival and construct design

To control protein trafficking to the plasma membrane, we used the previously established CAD-system that enables synchronizing secretory protein trafficking [13]. The construct (pCMV-CAD4-FCS-HA-POI-mCherry, Fig. 1a) consists of four CADs that retain the POI in the endoplasmic reticulum (ER) after synthesis by aggregate formation. Addition of a mixture of the small molecule D/D-Solubilizer and cycloheximide dissolves the aggregates and prohibits further protein synthesis, which leads to the release of the POIs from the ER in a synchronized wave. Later, the protease furin, which is located at the trans-Golgi network, cleaves at the furin cleavage site (FCS) and POIs without their CAD-tags are trafficked to the plasma membrane. After insertion into the plasma membrane, they expose the HA-tag to the extracellular space. This system has been characterized before for measuring the cell surface concentration of proteins with extracellularly applied antibodies (Fig. 1a,b) [5,13]. Here, we added a split luciferase fragment to the construct (pCMV-CAD4-FCS-Split_Luc-HA-POI-mCherry) that is also extracellularly exposed after protein insertion into the plasma membrane. The respective other split luciferase fragment was produced recombinantly in *E. coli* and added extracellularly. Thereby, the arrival of the proteins at the plasma membrane could be detected in real time by recombination of both fragments. Mardin Darby canine kidney (MDCK) cells were used because they readily form epithelial monolayers [3,14,15]. Rhodopsin (rhod) [13] and p75 [16] were used as markers for apical protein trafficking and the neural cell adhesion molecule (NCAM) [17] as basolateral marker.

**Figure 1.**
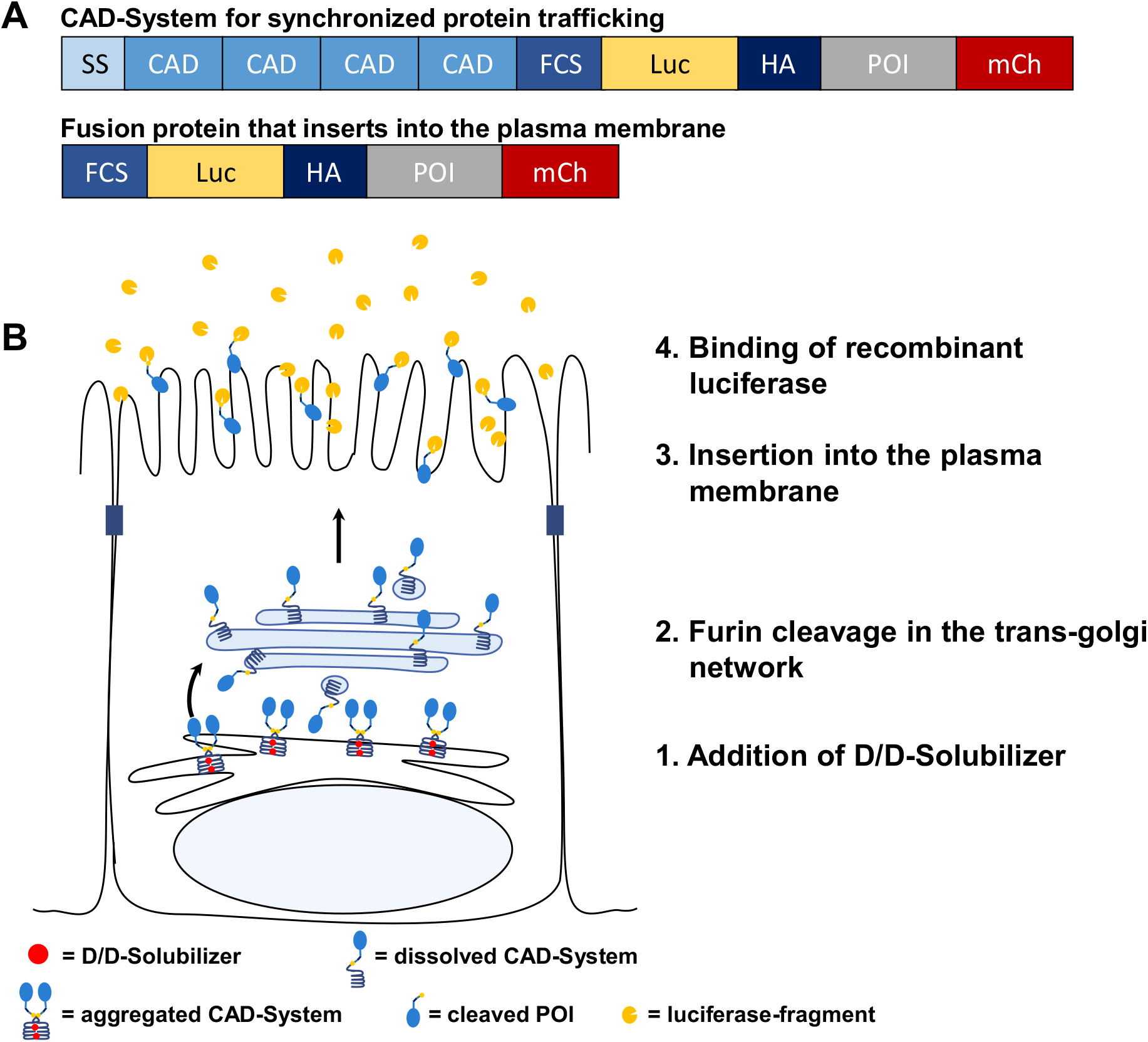
Construct design and mode of function of the split luciferase real-time reporter for protein plasma membrane arrival. (**A**) Construct for synchronized protein trafficking (“CAD-System”) consisting of a signal sequence (SS) for ER targeting, four conditional aggregation domains (CAD), a furin cleavage site (FCS), a split luciferase fragment (Split_Luc), HA-tag, the protein of interest (POI) and mCherry. POIs lacking the four CADs (lower construct) insert into the plasma membrane independent of D/D-Solubilizer addition. (**B**) POIs are first retained in the ER due to aggregation of the CADs. Addition of the D/D-Solubilizer dissolves these aggregates, and POIs traffic to the trans-Golgi network where the protease furin is located. Furin cleaves at the FCS and POIs without CAD tags progress to the plasma membrane. After insertion of the POI into the plasma membrane, the fused Split_Luc fragment and the HA-tag are exposed to the extracellular space and bind the recombinantly produced second Split_Luc fragment.

### System development and characterization

In our initial approach, we tested the detection of proteins arriving at the cell surface using the Gaussia luciferase and one of its mutants called “Monsta”. We split Gaussia luciferase between Gly93 and Glu94 and named them NGluc and CGluc, respectively [11]. Then, we inserted four point mutations in the N-terminus to create the Monsta variant, which is brighter and exhibits glow-type luminescence in comparison to the flash-type kinetics of wild type Gaussia luciferase [18]. We attached a hexa-histidine-tag (6x-His) either to the N-terminus or the C-terminus of the luciferase fragments and produced them in *E. coli*. The purified fragments were tested and the most promising variants were selected based on low background of the single fragments and high fold-induction after combination (Fig S1a). Combination of NGluc and CGluc led to significantly increased luminescence values and exchange of NGluc by Monsta further increased the signal 2.4-fold. Single NGluc and Monsta did not have activity alone, whereas CGluc had low intrinsic activity at high concentrations (Fig. 2a). We assumed that this activity was due to homooligomerisation of CGluc (Fig. S1b). As expected, wild-type Gaussia exhibited flash-type kinetics, whereas Monsta had higher and prolonged kinetics (Fig. 2b). Due to the low yield of CGluc production (Fig. S1b), and to avoid high background signal due to CGluc oligomers, we decided to fuse CGluc to our POIs (CGluc-HA-POI-mCherry) and produced stable cell lines that express CGluc-POI at the plasma membrane. We could only detect small quantities of CGluc-p75 and CGluc-NCAM at the plasma membrane, and CGluc-rhod was not detectable at all (Fig. S1c). We concluded that neither wild-type nor Monsta GLuc was sensitive enough to detect low protein amounts at the plasma membrane. Furthermore, the CGluc-tag impaired the efficiency of POI trafficking to the plasma membrane. This can be seen by comparing the microscopy images before and after CAD release from the ER (Fig. S2). In all three cases, a considerable amount of signal from the POI can be seen intracellularly, which indicates a problem with proper trafficking to the plasma membrane.

**Figure 2.**
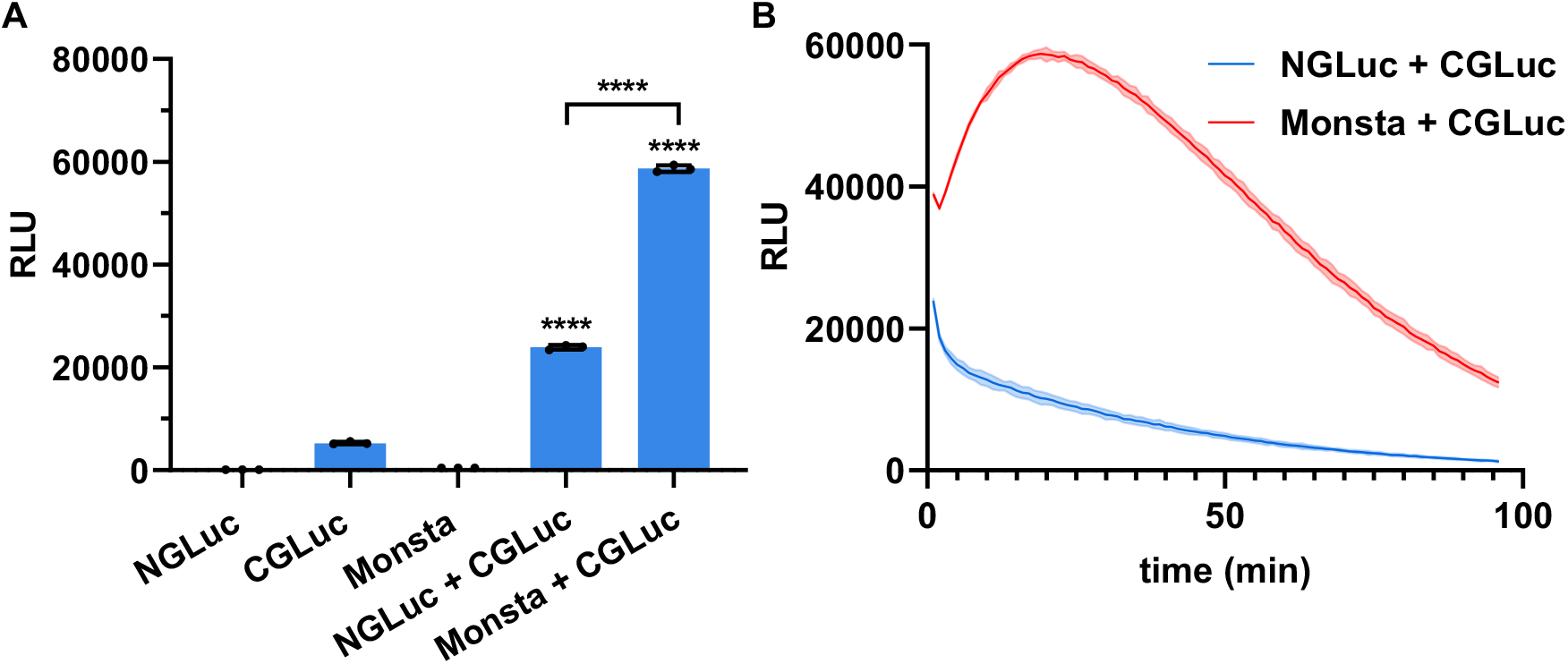
Activity of recombinantly produced split Gaussia luciferase fragments. (**A**) Indicated GLuc fragments were combined in 1:1 ratio at 1.25 μM concentration and luminescence in relative light units (RLU) was detected, maximum values of each time course are plotted, \*\*\*\**P* ≤ 0.001. (**B**) Distinct luminous behavior of wild-type GLuc and Monsta mutant over time. Fragments were combined in 1:1 ratio at 1.25 μM concentration, signals were measured every minute.

Therefore, we decided to test the NanoBiTs® system based on the Nanoluciferase for our purposes. Nanoluciferase is smaller, brighter and more stable compared to other luciferases and its split version is dissected asymmetrically into an 18 kDa, N-terminal part (LgBiT) and a 1.3 kDa, C-terminal part (SmBiT). The SmBiT is available with different affinities to the LgBiT. Therefore, we chose the SmBiT that was optimized for protein-protein interaction studies (SmBiT(114), K_D_ = 1.9 × 10^−4^) and two high affinity variants: SmBiT(86) (K_D_ = 0.7 × 10^−9^) and SmBiT(78) (K_D_ = 3.4 × 10^−9^) [12]. Due to its small size, SmBiT was chosen for fusion to our POIs rhod and NCAM. LgBiT was successfully produced in *E. coli* (Fig. S3). Analysis of POI localization by fluorescence microscopy before and after CAD release showed that SmBiT-tagged rhod and NCAM where not retained in the cytoplasm (Fig. S4).

In the next step, we tested whether we could detect the three SmBiT variants at the cell surface. To do so, we used constructs without the CADs (SmBiT-HA-POI-mCherry), in order for the POIs to be constantly present at the plasma membrane without the addition of the D/D-Solubilizer. We could detect both rhod and NCAM using the low-affinity SmBiT(114) (Fig. 3a,b). Luminescence signals only slightly increased 2.8-fold and 1.9-fold, respectively for rhod and NCAM, compared to the LgBiT background signal. Using the high affinity variants SmBiT(86) and SmBiT(78), fold-changes dramatically increased up to 6303-fold for rhod and 3441-fold for NCAM (Fig. 3a,b). This high sensitivity and dynamic range of the system opens the possibility to detect even low protein amounts at the plasma membrane and enables luminescence detection at the single cell level with a microscope (Fig. 3c).

**Figure 3.**
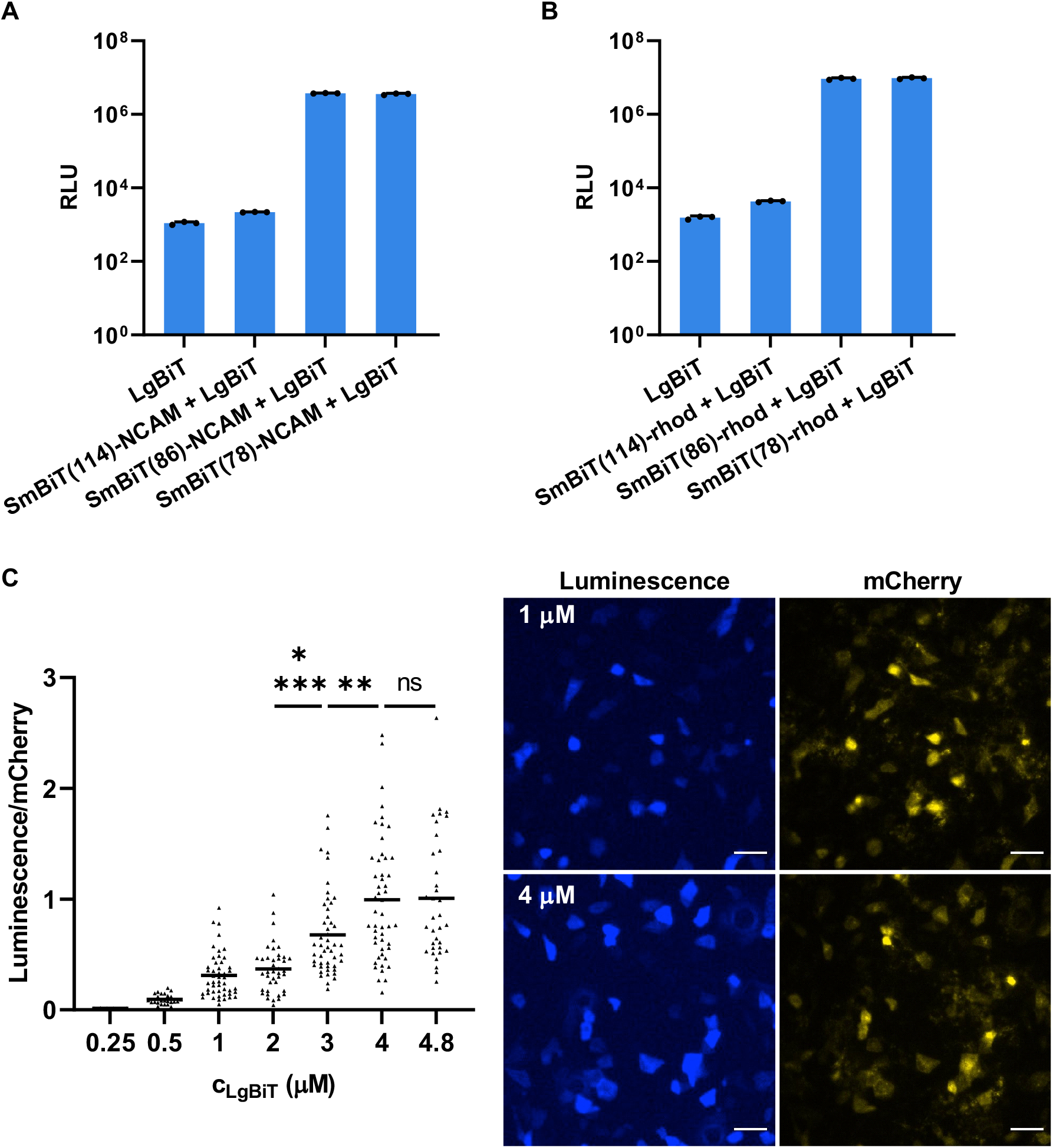
Detection of NCAM and rhod at the plasma membrane using split Nanoluciferase. (**A**,**B**) NCAM or rhod fused to the indicated SmBiT variant were transfected into unpolarized MDCK cells 12 h prior to the measurement. LgBiT was added at 83.3 nM concentration and substrate was used in 1:300 dilution. LgBiT in the extracellular medium showed a low intrinsic background activity, which is even detectable in untransfected cells. Mean values ± SD are plotted. (**C**) Detection of rhod at the cell surface via microscopy. Unpolarized MDCK cells were transfected with SmBiT(86)-rhod and LgBiT was added at the indicated concentrations. Luminescence and mCherry images were acquired with a confocal microscope using a 20x objective, scale bars indicate 50 μm. For quantitative analysis (left) background signals were subtracted and luminescence values were normalized to the mCherry signal of each cell. An outlier analysis was performed using GraphPad Prism (ROUT method, Q = 0.1%). Means with individual values are plotted (n.s., *P* ≥ 0.05; \*\**P* ≤ 0.01; \*\*\*\**P* ≤ 0.001), images of representative cells are shown (right).

### Real-time detection of plasma membrane arrival

After successful detection of SmBiT-tagged proteins that were stably expressed at the plasma membrane, we moved on to measure the dynamic arrival of trafficked proteins at the cell surface. To do so, we transiently expressed the CAD constructs for rhod and NCAM in polarized MDCK cells and added the D/D-Solubilizer simultaneously with the LgBiT and substrate to the cells. Luminescence signals were then measured every minute in a plate reader. We could detect the arrival of the first proteins after ≈ 30 min followed by the wave of the other synchronously trafficked proteins. The signal saturated after ≈ 120 min, indicating that all proteins had arrived at the plasma membrane (Fig. 4, Fig. S5a). SmBiT(78) worked in a similar manner, whereas for SmBiT(114) signal changes were barely detectable, pointing out the importance of a high affinity between LgBiT and SmBiT for spontaneous interaction and thereby providing a highly sensitive means for the detection of protein surface arrival (Fig. S5a,b).

**Figure 4.**
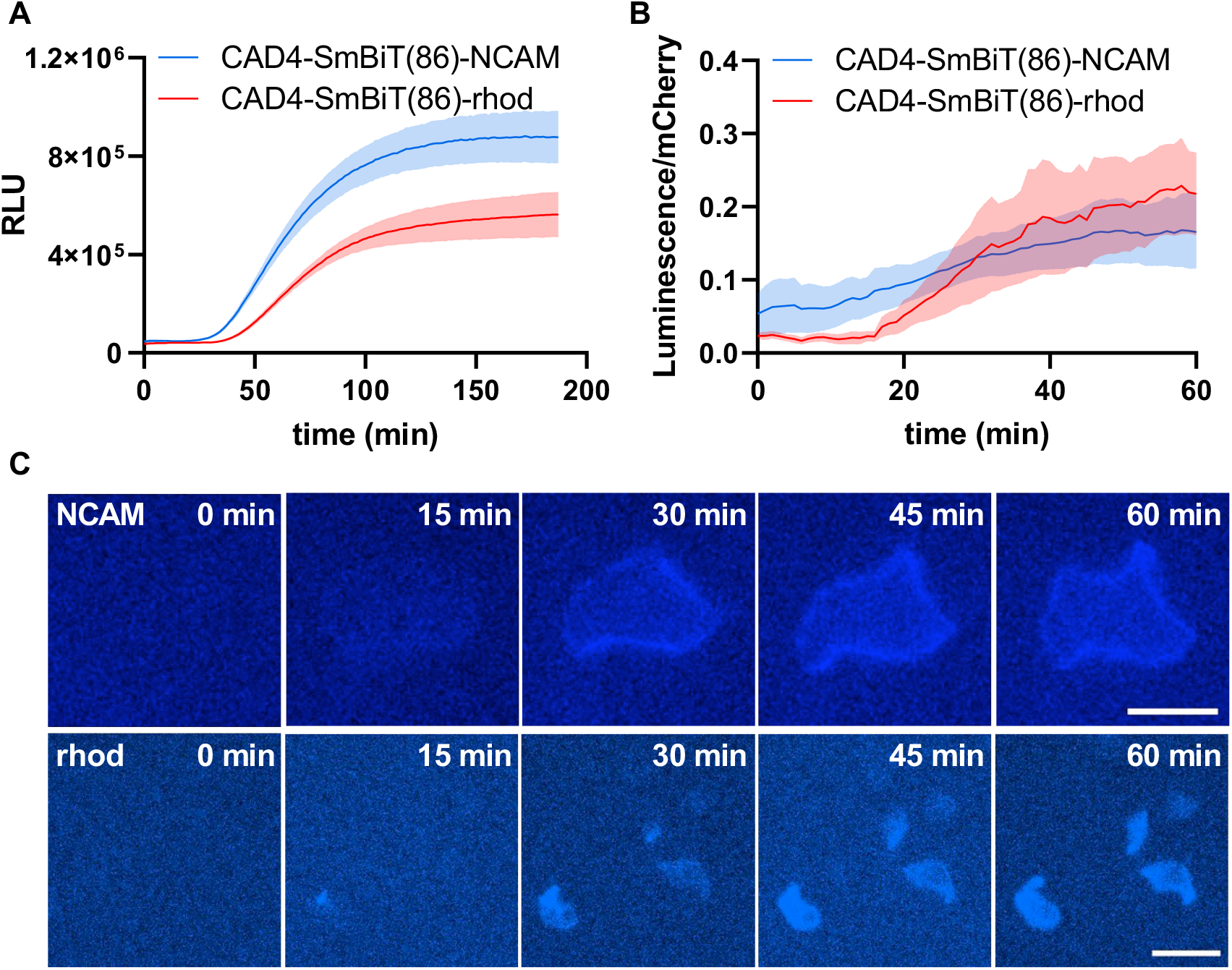
Real-time detection of protein plasma membrane arrival. (**A**) Plate reader measurements of unpolarized MDCK cells that were transfected with the indicated construct 12 h prior to the experiment. LgBiT was added at 83.3 nM concentration and substrate was used in 1:300 dilution. Mean values of each time point ± SD are plotted. (**B**) Quantitative analysis of real-time microcopy data of the plasma membrane arrival of NCAM (N=5) and rhod (N=12). Mean values of each time point ± SEM are plotted. (**C**) Representative images of selected time points for NCAM or rhod arrival at the plasma membrane. Images were acquired either with a 60x (NCAM, scale bar: 20 μm) or a 20x (rhod, scale bar 50 μm) objective.

A typical problem with long luciferase measurements is the loss of substrate activity because of instability or degradation of enzymes over time. To distinguish whether signal decay after reaching the plateau derived from an actual change of protein amount, for example due to protein internalization, or simply from loss of substrate activity, we made control measurements for each experiment (Fig. S5c). To control for these effects, we expressed the respective constructs without the CADs and fixed the cells. In these controls, signal changes should have occurred exclusively due to changes in substrate activity. We used this data to normalize our protein arrival curves (Fig. S5d).

Next, we confirmed our plate reader data by monitoring protein arrival at the plasma membrane in real time by microscopy. We recorded time series over 60 min and could show the arrival of rhod and NCAM at the plasma membrane (Fig. 4b). The spatial resolution was high enough to see a characteristic plasma membrane signal for NCAM, which corresponds to the signal distribution that is expected to be seen in a microscopic z-section of a basolateral protein like NCAM (Fig. 4c). These recordings open up entirely new possibilities for the spatial and temporal detection of protein plasma membrane arrival at single cell level.

### Specific detection of apical and basolateral protein arrival

Finally, we wanted to test if our system is capable of specifically distinguishing protein arrival at the apical and basolateral plasma membrane. Therefore, we grew MDCK cells on the bottom of transwell filters until they formed a continuous polarized monolayer and transfected them with the constructs without the CADs. LgBiT and substrate was then added either to the apical or the basolateral side of the cell layers for polarity-specific detection (Figure S6). For NCAM, signals were only detectable if substrate and LgBiT were added to the basolateral side of the cells (Fig. 5a) and for rhod it was *vice versa* (Fig. 5b). With this experiment we could show that our system based on split Nanoluciferase is able to detect the cell surface polarity of apical and basolateral membrane proteins precisely and quantitatively.

**Figure 5.**
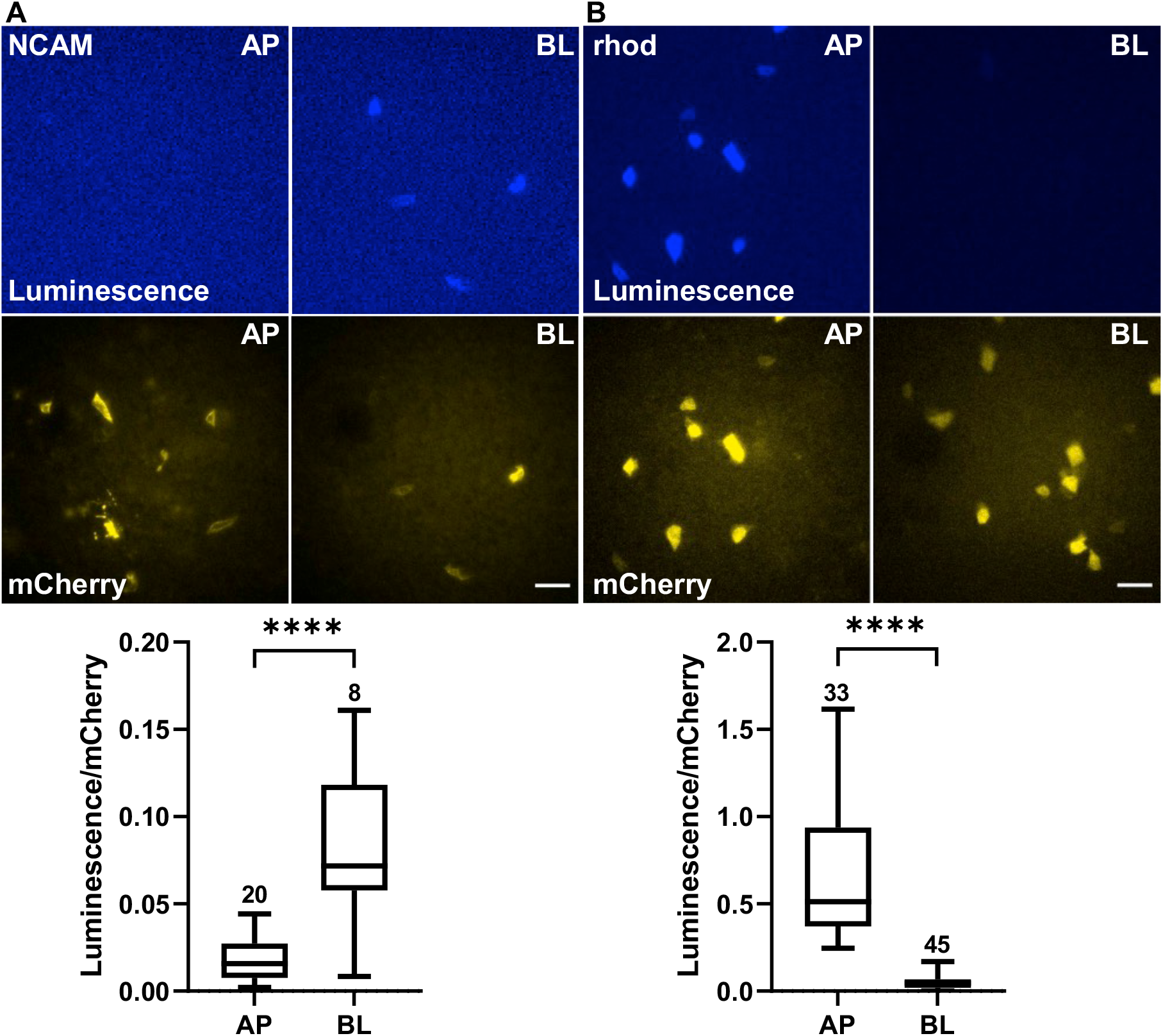
Apico-basal polarity-specific detection of NCAM and rhod. (**A**,**B**) Representative images and quantitative analysis of MDCK cells expressing SmBiT(86)-NCAM or SmBiT(86)-rhod constitutively at the plasma membrane. Cells were grown on the bottom of transwell filters until they formed a polarized monolayer. 3 μM LgBiT and substrate diluted 1:100 in PBS was added apically or basolaterally directly prior measurement. NCAM images were acquired with 100 ms exposure time for mCherry and 2 sec for luminescence. For rhod, 50 ms exposure time was used for mCherry and 1 sec for luminescence, scale bars indicate 50 μm. For analysis, background signals were subtracted and the luminescence signal was normalized with mCherry signal of each cell. N is indicated above each boxplot, \*\*\*\**P* ≤ 0.0001.

## Discussion

Maintaining the concentration of cell surface proteins within tight boundaries is crucial for the proper function of a cell within a multicellular organism. However, directly measuring the cell surface concentration of a particular protein of interest (POI) in live cells in real time represents a considerable challenge. Here, we introduce an approach based on split luciferases and characterize it by biochemical and cell biological methods and demonstrate its applicability by using it to measure the cell surface arrival of CAD-synchronized proteins.

### Comparison of the utility of split Gaussia luciferase and split Nanoluciferase for real-time protein cell surface arrival assays

In our initial trials, we used split Gaussia luciferase including its ‘Monsta’ variant. However, our experiments showed that split Gaussia luciferase does not have favorable properties for using it in split luciferase cell surface arrival assays. First, the signals after recombining split Gaussia luciferase were not high enough to enable sensitive measurements on a single cell level. Second, a rather large fragment of split Gaussia luciferase needed to be attached to the POI, which often resulted in side effects on the natural trafficking of the POI. In comparison, split Nanoluciferase showed much higher signals after recombination. We measured a more than 6000-fold signal increase above background. Furthermore, only a 11 amino acid fragment needs to be attached to the POI, which makes interference with the natural function of the POI much less likely. Nevertheless, the purified large fragment of split Nanoluciferase showed slight luminescence activity even without the presence of the small fragment. This could be due to other small peptides present in the extracellular medium, but the exact reason for this remains unknown.

### Applicability of our approach for studying polarized membrane trafficking in epithelial cells

We showed that our method can be used to monitor the polar cell surface arrival of apical and basolateral plasma membrane proteins in polarized epithelial cells. As we also demonstrated, the sensitivity of our system is sufficient to use it for single-cell experiments with a microscopic readout. This opens a multitude of novel possibilities. Classic methods to characterize protein sorting and polarized cell surface arrival in epithelial cells are based on biochemical readouts and utilize immunoprecipitation to measure radioactively pulse-chase labelled proteins and/or proteins that are labelled at the cell surface with reactive NHS-esters [17,19]. Although these biochemical methods show a very high sensitivity, they are not able to reveal data on a single-cell level. Alternatively, POIs at the cell surface can be labeled with extracellular antibodies, which allows to read out the cell surface concentration of POIs at the individual cell level. However, antibody-based methods cannot be applied for live cell real-time measurements, because unbound antibodies need to be washed out. We carried out such antibody-based assays in previous studies [5,13] and clearly saw that cell surface arrival varies significantly between individual cells.

In summary, virtually all data available on proteins or mutations that are relevant for polarized apical or basolateral protein sorting, are end-point-measurements and/or bulk averages, and therefore might hide or ‘average out’ several intricacies and/or sub-states during the sorting process. With the new split luciferase-based approach presented here, it will become possible to characterize and study these variations in single cells over time. It will be an interesting future endeavor to correlate the observed trafficking variations with variations in the expression and activity of sorting protein complexes at a single cell level. The split luciferase-based approach here would even make it possible to carry out such studies in more complex model systems such as organ explants or organoids.

## Materials and Methods

### DNA cloning and production

Plasmids were cloned by restriction enzyme cloning or Gibson assembly [20]. Short DNA fragments up to 80 bp were inserted by aligning two compatible primers (Merck KGaA) that mimic specific restriction enzyme sites at both ends. The alignment was achieved by mixing 20 μL of each forward and reverse primer (100 μM) with 20 μL 20 M NaCl-solution and 20 μL H_2_O and heating to 95 °C for 10 min. The mix was slowly cooled down to room temperature, diluted 1:100 and used for the ligation using the Quick Ligation Kit (New England Biolabs). Specific base pair substitutions were introduced with site directed mutagenesis by polymerase chain reaction (PCR). Sequences were verified by Sanger sequencing. All used plasmids are listed in Supplementary table 1.

### Recombinant protein production and purification

Recombinant luciferase fragments were produced in *E. coli* BL21 (DE3). 1 L bacterial cultures were grown at 37°C, 150 rpm in LB-Medium (Roth). Upon reaching OD_600_ = 0.6 - 0.8, the expression was induced with 1 mM isopropyl β-D-1-thiogalactopyranoside (IPTG). The expression was sustained for 20-24 h at 18 °C and 150 rpm. Cultures were harvested by centrifugation (6000 *g*, 10 min) and pellets were resuspended in 30 mL nickel (Ni)-lysis buffer (50 mM NaH_2_PO_4_ x 2 H_2_O, 300 mM NaCl, 1 mM Imidazole, pH 8). Lysates were shock frozen in liquid nitrogen and stored at -80 °C. Recombinant proteins were purified over nickel-nitrilotriacetic acid (Ni-NTA) Agarose (Qiagen). Therefore, lysates were thawed in a water bath at 37 °C, sonicated (Bandelin Sonoplus HD 3100 homogenizer) for 10 min at 60 % amplitude on ice and centrifuged for 1 h with 30.000 *g* at 4 °C. Polypropylene gravity-flow columns (Thermo Fisher) were prepared with 1 mL Ni-NTA Agarose according to the manufacturers protocol and equilibrated twice with 15 mL Ni-lysis buffer. Cleared lysates were added to the columns and subsequently washed twice with 15 mL Ni-wash buffer (50 mM NaH_2_PO_4_ x 2 H_2_O, 300 mM NaCl, 20 mM Imidazole, pH 8). Elution with Ni-Elution Buffer (50 mM NaH_2_PO_4_ x 2 H_2_O, 300 mM NaCl, 250 mM Imidazole, pH 8) was done in 1 mL steps. Protein content was confirmed by mixing 2 μL of each elution step with 200 μL Bradford solution (Protein assay dye reagent (Bio-Rad) diluted 1:5 in H_2_O). Eluates containing protein were pooled and analyzed via SDS-PAGE and coomassie brilliant blue staining. Protein concentration was determined via Bradford protein assay and adjusted to 0.1 mg/mL in phosphate buffered saline (PBS) containing calcium, magnesium (PBS (+/+), Life Technologies) and 0,1 % glycerol to be stored in aliquots at -80 °C.

### Mammalian cell culture and transfection

Madin-Darby Canine Kidney cells subclone II (MDCK II, kindly provided by Enrique Rodriguez-Boulan, Weill Cornell Medical College) were cultivated at 37 °C and 5 % CO_2_ in Dulbecco’s Modified Eagle Medium (DMEM, Life Technologies) containing 5 % fetal calf serum (FCS, Life Technologies) and 2 mM L-glutamine. Cells were passaged after reaching a confluency of 80 – 90 %. For experiments, cells were seeded onto assay plates and transfected using polyethyleneimine (PEI, Polyscience, 1 mg/mL). Details on the conditions for each experiment can be found in Supplementary Table 2. For “synchronized release” experiments MDCK II cells were grown in a polarized and confluent monolayer. Therefore, cells were seeded on transwell filters (#3401, Corning-Costar) and grown for 4 days. Medium was changed daily. 8 h after transfection, cells were incubated with D/D-Solubilizer (5 μM, Clonetech/Takara) and Cycloheximide (35.54 μM, Merck) for 3 h at 37 °C.

### Generation of stable cell lines

To generate stable cell lines, 2.5 × 10^5^ cells were seeded on a 6-well plate. The next day they were transfected with the desired vector, containing a resistance to geneticin (G418). After overnight incubation, cells were detached from the plate, diluted 1:10, 1:20, 1:50 and 1:200 with DMEM containing 5 μg/mL G418 (Roth) and seeded on 10 cm dishes. Cells were then grown for approximately 2 weeks until single colonies were grown. The medium was refreshed twice a week. Single colonies were isolated and mCherry positive colonies were selected by confocal microscopy.

### Luciferase readout

Luciferase experiments in 96-well format were measured using a Tecan Infinite M200 Pro which measures luminescence signals in a spectrum from 380 – 600 nm. All measurements were performed in triplicates in white, flat-bottomed 96-well plates with an integration time of 1000 ms. Substrate was always prepared freshly, protected from light and added directly prior the measurement. For the Gaussia luciferase, 100 μL injections of coelenterazine (Roth, 20 μM, in PBS) were used and for the Nanoluciferase, 50 μL injections of Furimazine (Nano-Glo® Luciferase Assay Substrate, Promega) were applied. The substrate was diluted 1:50 in Nano-Glo® Luciferase Assay Buffer for recombinant proteins or in PBS for luminescence measurement in cells. Recombinant luciferase fragments were always thawed freshly for each experiment, combined in the desired concentrations and measured at room temperature. To detect luciferase fragments on the surface of cells, stable cell lines or cells to be transfected 24 h prior the measurement were seeded onto tissue culture coated, white 96-well plates 48 h prior the experiment. If the steady amount of luciferase fragments at the surface was measured, cells were fixed with 4 % paraformaldehyde solution (PFA) for 15 min at room temperature, washed with PBS and remaining PFA was quenched with 50 mM NH_4_Cl for 2 min at room temperature. Afterwards, the respective other recombinant fragment of the split luciferase was added in the desired concentration.

To detect protein arrival at the cell surface, cells were washed with PBS and respective recombinant luciferase fragment was added. D/D-Solubilizer (5 μM) was mixed with the substrate and added directly prior the measurement. Protein surface arrival was measured at 37 °C, the arrival curve was corrected for the loss of substrate activity by normalization to the control samples (Fig S5).

### Sample preparation for Microscopy

To detect HA-tagged proteins at the cell surface, a HA-Cy5-antibody (mouse anti-HA antibody (Covance, PRB-101C), conjugated to Cy5 using a Cy5 conjugation kit (GE Healthcare) was diluted 1:500 in medium and applied to the apical and basolateral side of the transwell filters. After 15 min incubation at 37 °C, cells were washed twice with PBS, fixed with 4 % PFA for 15 min at room temperature and washed again with PBS. Staining of nuclei was performed by incubating the samples for 30 min with 1 μg/mL DAPI (Thermo Fisher Scientific) in SAPO solution (0.02 % saponin, 0.2 % BSA in PBS). Samples were washed three times with PBS. Coverslips were mounted in 8.5 μL Mowiol mounting medium (200 % Tris HCl (0.2 mM, pH 8.5), 100 % Glycerol, 40 % Mowiol (Roth) in H_2_O), supplied with 1,4-Diazabicyclo[2.2.2]octan-medium (Dabco-medium, (Tris HCL (50 mM, pH 8), 2.5 % Dabco (Merck) in glycerol), whereas transwell filters were excised and mounted in Dabco-medium. Live cell imaging was performed in live cell imaging chambers (Ibidi) or on transwell filters. To enable imaging with transwell filters, the cells were seeded onto the downside of transwell filters. To do so, the transwell inserts were placed bottom-up on a 10 cm dish and 400 μL cell suspension was added. The dish was then carefully covered with the lid, without destroying the drop of cell suspension and incubated for 3 h until the cells attached to the filter. Then the filters were transferred back into a 12-well plate and covered with medium. In order to grow a tight and intact monolayer, the medium was changed every day for 7 days. For transfection, the transwell filters were again placed bottom up on a 10 cm dish, the transfection mix was applied and incubated for 6-8 h at 37 °C. The filters were then transferred back into the 12-well plate and incubated overnight. Directly before imaging, the filters were carefully placed on a 32 mm live cell imaging chamber with 750 μL PBS or DMEM without phenol red (Fig. S6). For the experiment, PBS or DMEM was replaced by the respective recombinant luciferase fragment that was mixed with the substrate and added either apically or basolaterally.

### Imaging and image analysis

Fluorescent images were acquired using a Nikon Eclipse Ti.E A1R inverted confocal laser scanning microscope equipped with a 60X oil immersion objective (numerical aperture (N.A.) = 1.49) and lasers emitting at 405, 488, 561, and 641 nm. Z-stacks covering the complete cell height were recorded and cells were analyzed using a custom-written Matlab program [5,13,21]. Luciferase signals were imaged by combining the A1R confocal setup with an ANDOR DU-897 EMCCD camera and a 20x plan apochromatic objective (N.A. = 0.75). To detect the full luminescence spectrum, no filter cube was used. For image analysis, the background was subtracted and luminescence signals were normalized to mCherry signals. For time series, several positions on the sample were chosen, and one picture per minute was recorded over a period of 2-3 h.

## Data Availability

Data are available upon request from.

## Acknowledgements

RT acknowledges support from the Ministry of Science, Research and the Arts of Baden-Württemberg (project: Split luciferase-mediated detection of trafficked protein localization). This work was further supported by the German Research Foundation (Deutsche Forschungsgemeinschaft, DFG) under Germany’s Excellence Strategy – CIBSS, EXC-2189, Project ID: 390939984 and under the Excellence Initiative of the German Federal and State Governments – BIOSS, EXC-294 and SGBM, GSC-4, the grant RO 4341/2-1 and in part by the Ministry for Science, Research and Arts of the State of Baden-Württemberg (Az: 33-7532.20).

