## Supplementary Information for "Real-time monitoring of cell surface protein arrival with split luciferases"

**Running title:**

Luciferases to measure surface arrival

Alexandra A.M. Fischer[1,2,3], Julia Baaske[1,2], Winfried Römer[1,2], Wilfried Weber[1,2,3], Roland Thuenauer[4,5,6,*]

[1] Signaling Research Centres BIOSS and CIBSS and Faculty of Biology, University of Freiburg, Freiburg, Germany

[2] Faculty of Biology, University of Freiburg, 79104 Freiburg, Germany

[3] Spemann Graduate School of Biology and Medicine (SGBM), University of Freiburg

[4] Center for Structural Systems Biology (CSSB), Hamburg, Germany

[5] Technology Platform Light Microscopy, University of Hamburg, Hamburg, Germany

[6] Technology Platform Microscopy and Image Analysis (TP MIA), Leibniz Institute of Virology (LIV), Hamburg, Germany

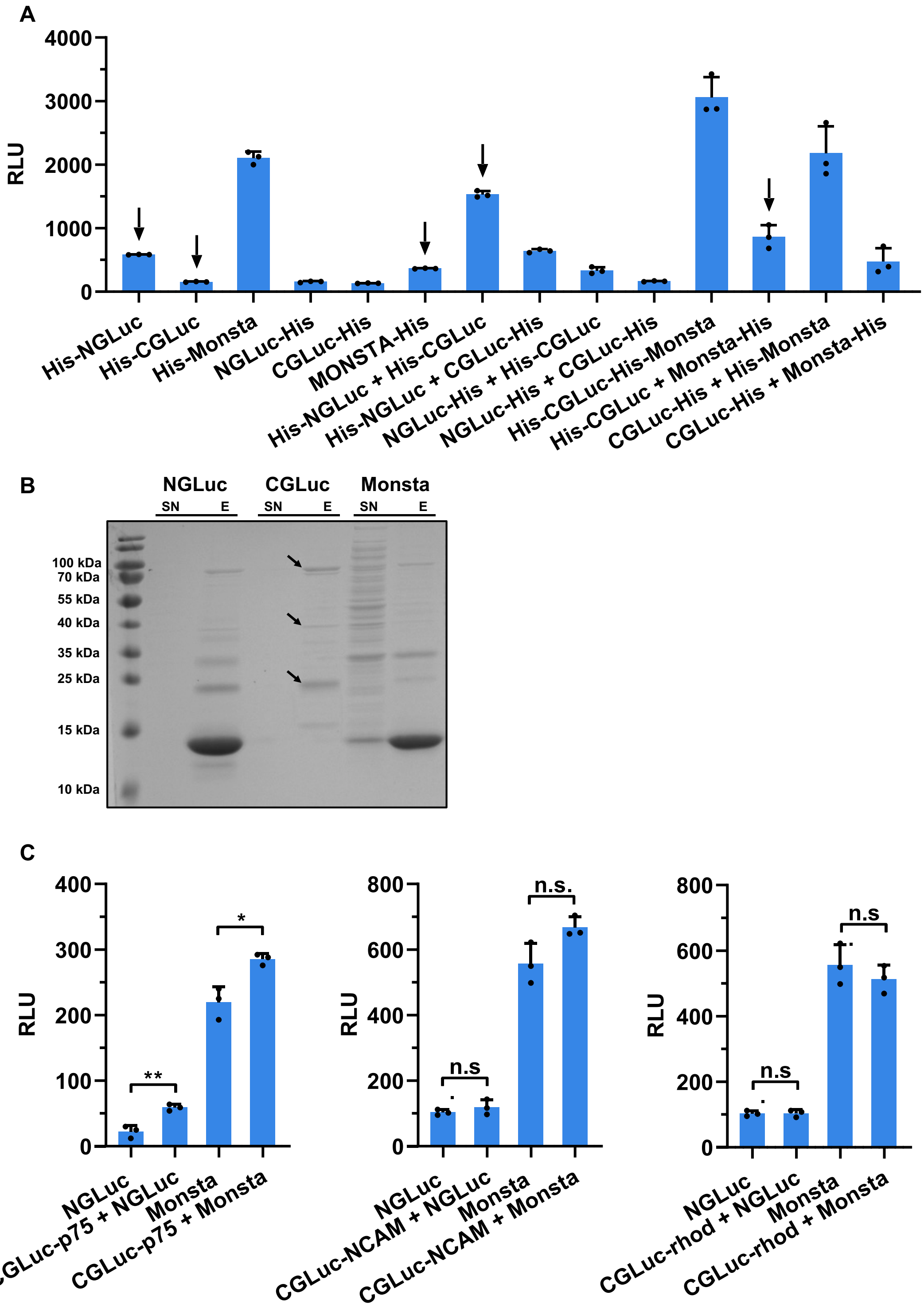


**Figure S1**. (**A**) Comparison of all recombinant GLuc variants. Proteins were combined in 1:1 ratio at 2.5 μM concentration, mean luminescence values ± SD are plotted. Arrows indicate the variants that were selected for further experiments. (**B**) Coomassie brilliant blue stained SDS gel of a large-scale purification of the GLuc fragments. NGLuc and MONSTA with His(6x)-tag have a molecular weight of 12 kDa, His-CGLuc has 10 kDa but seems to form multimers that are indicated with the arrows. Supernatant (SN) and eluate (E) of the purification were loaded. (**C)** NGLuc or Monsta were added at 1.25 μM concentration to stable cell lines expressing CGLuc-p75, CGLuc-NCAM or CGLuc-rhod without CADs constitutively at the cell surface (n.s., $P \geq 0.05$; $*P \leq 0.05$; $**P \leq 0.01$).

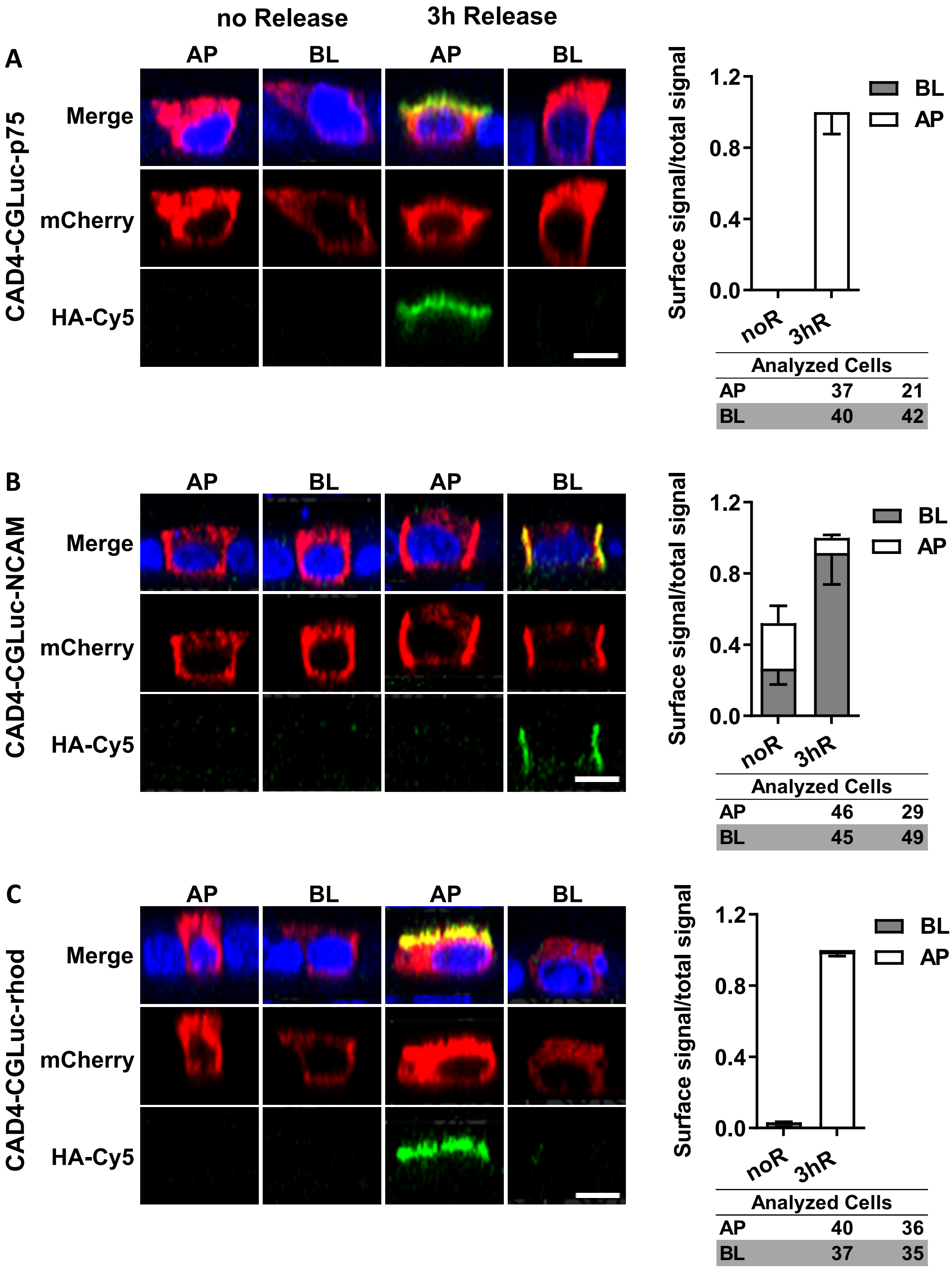


**Figure S2**. Quantitative analysis of the synchronized release of CGLuc-tagged p75 (**A**), NCAM (**B**) and Rhod (**C**). MDCK cells, grown as a polarized monolayer, expressed the indicated constructs. They were incubated for 3 h (release) with or without (no release) D/D-Solubilizer. Afterwards, HA-Cy5-antibody surface staining was performed apically (AP) or basolaterally (BL). The ratio of the surface signal to total signal ± SEM is plotted, N is indicated underneath. Images show representative cells in xz or yz-view, the scale bar indicates 10 µm.

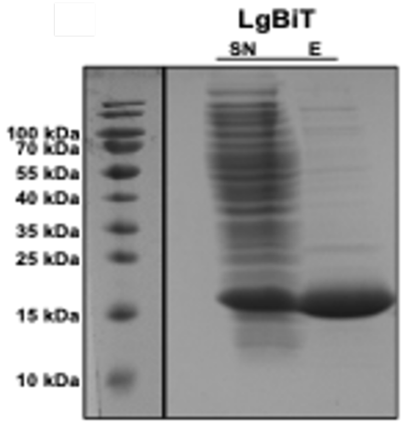


**Figure S3**. Representative coomassie brilliant blue stained SDS gel of supernatant (SN) and eluate (E) of the protein purification of (6x)-His-tagged LgBiT (20 kDa).

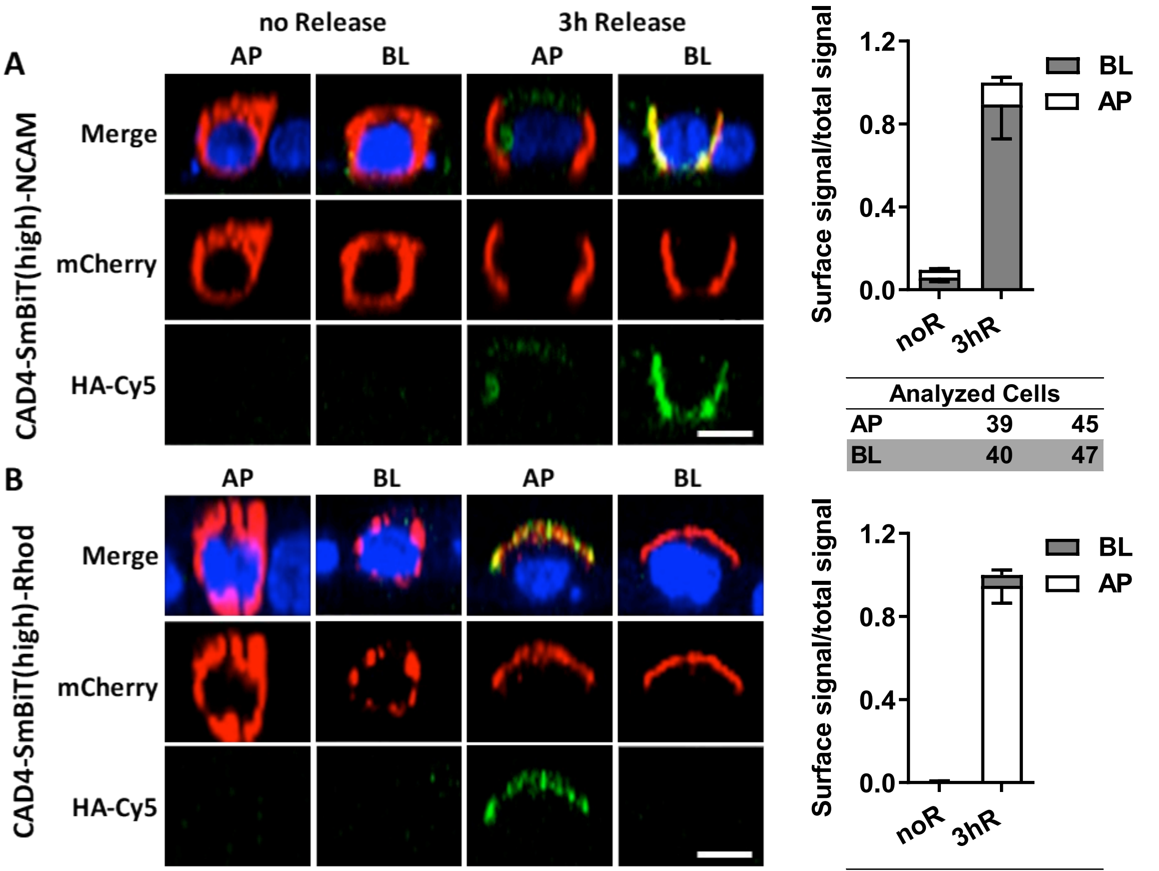


| Analyzed Cells | | |
|---|---|---|
| AP | 39 | 45 |
| BL | 40 | 47 |

| Analyzed Cells | | |
|---|---|---|
| AP | 49 | 43 |
| BL | 49 | 57 |

**Figure S4.** Quantitative analysis of the synchronized release of SmBiT(86)-tagged NCAM (**A**) and Rhod (**B**). MDCK cells, grown as a polarized monolayer, expressed the indicated constructs. They were incubated for 3 h (release) with or without (no release) D/D-Solubilizer. Afterwards, HA-Cy5-antibody surface staining was performed apically (AP) or basolaterally (BL). The ratio of the surface signal to total signal ± SEM is plotted, N is indicated in the tables underneath. Images show representative cells in xz- or yz-view, the scale bar indicates 10 µm.

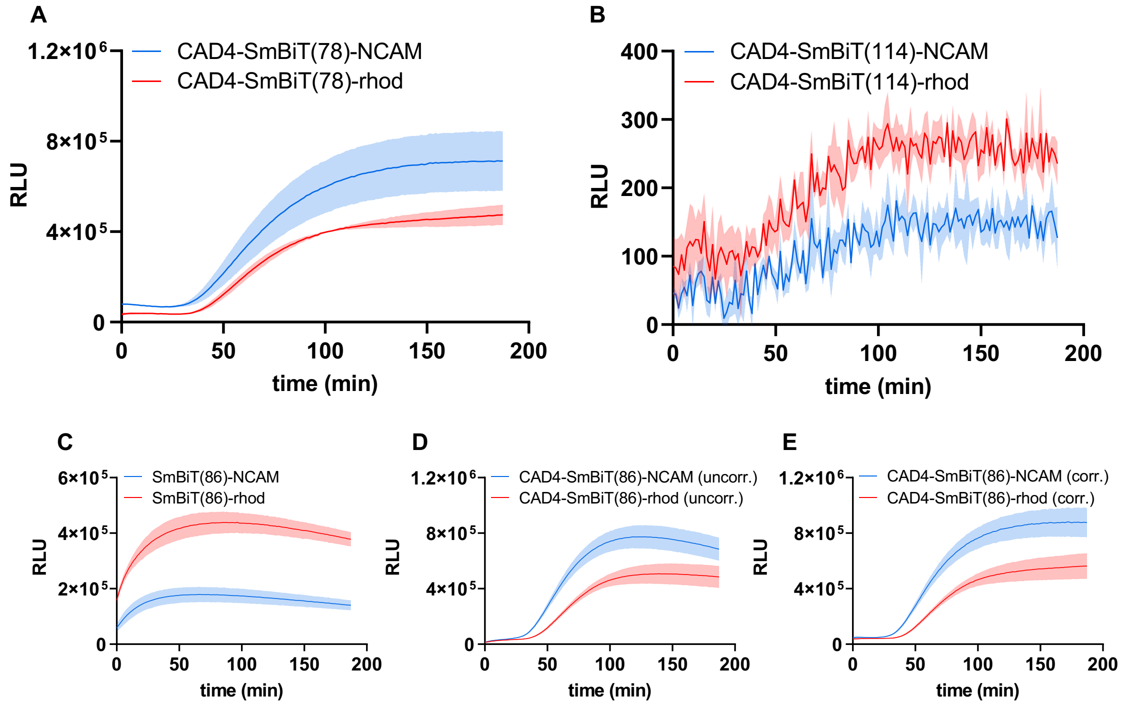


**Figure S5**. Real-time measurements of NCAM and rhod arrival at the plasma membrane. (**A, B**) Arrival kinetics using the SmBiT variants 78 (high LgBiT affinity) and 114 (low LgBiT affinity). (**C-E**) Curve correction procedure. (**C**) Measurement of control curves with constructs without the CADs that are constitutively expressed at the cell surface. (**D**) Uncorrected arrival curves for the indicated constructs. (**E**) Arrival of NCAM and rhod with correction for loss of substrate activity by normalization to the control curves and the maximum.

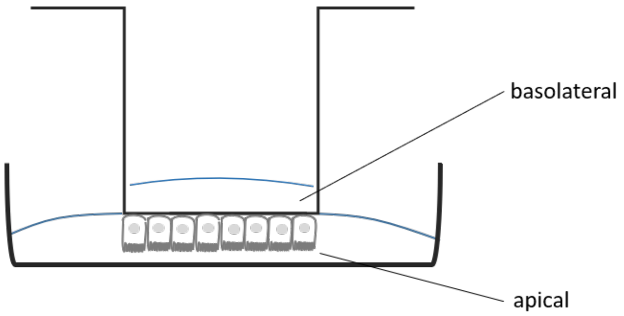


**Figure S6**. Scheme of the preparation of MDCK cells for live cell imaging. Cells were seeded and grown on the downside of transwell filters. For imaging they were placed in PBS or DMEM without phenol red and LgBiT and substrate were added either exclusively to the apical or to the basolateral side of the cells.

**Table S1:** Plasmids used in this study

| Name | Description | Backbone |
|---|---|---|
| pAF1003 | SS-CAD4-FCS-CGluc-HA-NCAM-mCh | pEGFP-N1 |
| pAF1006 | SS-CAD4-FCS-CGluc-HA-p75-mCh | pEGFP-N1 |
| pAF1009 | SS-CAD4-FCS-SmBiT(114)-HA-NCAM-mCh | pEGFP-N1 |
| pAF1013 | His(6x)-Thrombin cleavage site-NGLuc | pET15b |
| pAF1014 | His(6x)-Thrombin cleavage site-CGLuc | pET15b |
| pAF1015 | His(6x)-Thrombin cleavage site-NGLuc_MONSTA | pET15b |
| pAF1016 | His(6x)-Thrombin cleavage site-LgBiT | pET15b |
| pAF1018 | NGLuc-Thrombin cleavage site-His(6x) | pET15b |
| pAF1019 | CGLuc-Thrombin cleavage site-His(6x) | pET15b |
| pAF1020 | NGLuc_MONSTA-Thrombin cleavage site-His(6x) | pET15b |
| pAF1021 | LgBiT-Thrombin cleavage site-His(6x) | pET15b |
| pAF1024 | FCS-CGluc-HA-NCAM-mCh | pEGFP-N1 |
| pAF1027 | FCS-CGluc-HA-p75-mCh | pEGFP-N1 |
| pAF1030 | FCS-SmBiT(114)-HA-NCAM-mCh | pEGFP-N1 |
| pAF1032 | FCS-SmBiT-HA-p75-mCh | pEGFP-N1 |
| pAF1033 | SS-CAD4-FCS-CGLuc-HA-Rhod-mCh | pEGFP-N1 |
| pAF1034 | SS-CAD4-FCS-SmBiT(114)-HA-Rhod-mCh | pEGFP-N1 |
| pAF1035 | FCS-CGLuc-HA-Rhod-mCh | pEGFP-N1 |
| pAF1036 | FCS-SmBiT(114)-HA-Rhod-mCh | pEGFP-N1 |
| pAF1038 | SS-CAD4-FCS-SmBiT(86)-HA-NCAM-mCh | pEGFP-N1 |
| pAF1039 | SS-CAD4-FCS-SmBiT(86)-HA-Rhod-mCh | pEGFP-N1 |
| pAF1040 | SS-CAD4-FCS-SmBiT(78)-HA-NCAM-mCh | pEGFP-N1 |
| pAF1041 | SS-CAD4-FCS-SmBiT(78)-HA-Rhod-mCh | pEGFP-N1 |
| pAF1042 | FCS-SmBiT(86)-HA-NCAM-mCh | pEGFP-N1 |
| pAF1043 | FCS-SmBiT(86)-HA-Rhod-mCh | pEGFP-N1 |
| pAF1044 | FCS-SmBiT(78)-HA-NCAM-mCh | pEGFP-N1 |
| pAF1045 | FCS-SmBiT(78)-HA-Rhod-mCh | pEGFP-N1 |

**Table S2:** Cell numbers and transfection conditions

| Dish | Growth state | Cell Number | PEI (µL) | DNA (µg) | OptiMEM (µL) |
|---|---|---|---|---|---|
| 6-well | unpolarized | $2.5 \times 10^5$ | 3.6 | 11.25 | 900 |
| 12 mm transwell-filter | polarized | $2 \times 10^5$ | 5 | 1.6 | 200 |
| 24-well | unpolarized | $5 \times 10^4$ | 2.5 | 0.8 | 200 |
| 96-well | unpolarized | $1.2 \times 10^4$ | 0.781 | 0.25 | 30 |
| 8-well µ-Ibidi slide | polarized | $2.5 \times 10^5$ | 5 | 1.6 | 105 |
